# Estimating diversification rates from the fossil record

**DOI:** 10.1101/027599

**Authors:** Gilles Didier, Marine Fau, Michel Laurin

## Abstract

Diversification rates are estimated from phylogenies, typically without fossils, except in paleontological studies. By nature, rate estimations depend heavily on the time data provided in phylogenies, which are divergence times and (when used) fossil ages. Among these temporal data, fossil ages are by far the most precisely known (divergence times are inferences calibrated with fossils). We propose a method to compute the likelihood of a phylogenetic tree with fossils in which the only known time information is the fossil ages.

Testing our approach on simulated data shows that the maximum likelihood rate estimates from the phylogenetic tree shape and the fossil dates are almost as accurate as the ones obtained by taking into account all the data, including the divergence times. Moreover they are substantially more accurate than the estimates obtained only from the exact divergence times (without taking into account the fossil record).

## 1 Introduction

The history of biodiversity, including evolutionary radiations and mass extinction events, is a central topic of macroevolutionary studies (see for instance Santini *et al*., 2013; Sidor *et al*., 2013; Benton *et al*., 2014, and other works from these authors). Yet, despite the fact that the most direct data about these subjects is found in the fossil record, the latter is currently under-exploited in such studies because sophisticated methods that could assess cladogenetic (speciation) and extinction rates are still under development (Stadler, 2010; Didier *et al*., 2012). Paleontologists have long studied these phenomena, but with simple methods that typically account, at best, only for ghost lineages (Norell, 1993), but that otherwise assume a fairly literal reading of the fossil record.

In parallel, evolutionary biologists working on extant taxa have developed sophisticated methods to study the diversification of taxa, but usually, without incorporating data from the fossil record. For instance, Gernhard (2008) studied the diversification from a theoretical point of view and considered cases in which all the data were from extant taxa. This work has been generalized by Hallinan (2012), who provided several useful results, but who stated, erroneously, that “all real phylogenies end in the present”. In fact, many large clades are entirely extinct (Carroll, 1988), such as stereospondyls, dinocephalians dicynodonts, and ornithischian dinosaurs, to mention but a few, so their phylogenies end in the past, but it would be useful to develop similar methods to deal with such cases, as paleontologists have studied the evolutionary radiations of such taxa (Ruta *et al*., 2007, 2011). Some developments in that direction were made recently (Stadler, 2010; Didier *et al*., 2012). Below, we present further developments of such a method.

*Diversification* is an user-friendly software which estimates the diversification and fossilization rates from tree shapes and fossil dates. It has tree/fossil occurrence editing capabilities and it is available at GitHub, altogether with a console version and several utilities (https://github.com/gilles-didier/Diversification).

## 2 Diversification process, fossils and reconstructions

### 2.1 Reconstructing evolution from the extant taxa and the fossil record

A prerequisite for studying the diversification process is to extract whatever information can be obtained about it from the present day, which will be referred to as time *T*. Until we can travel through time, we have only access to the contemporary lineages and the fossil record. This means that much data about the evolutionary pattern are irremediably lost. In particular, all the lineages data about extinct without leaving any fossils are definitively out of reach.

Let us formalize what can be reconstructed all available data. A lineage *ℓ,* alive at time *t*, is said to be *observable at time t* if it does not go extinct before *T* or if a fossil of *ℓ* or of one of its descendants is found at a time greater than *t*. The part of the evolutionary tree (with fossil finds; for simplicity below, we refer to this as a “tree”, but it should be understood that this includes information about fossil finds, unless noted otherwise) that can be reconstructed is what is observable. Basically the observable part of the whole evolutionary tree is obtained by pruning the branches of extinct lineages just after the time of their most recent fossil finds, or removing them if they have no fossil record. The observable part of a tree will be referred to as the *reconstructed tree with fossils.*

### 2.2 Modeling both diversification and fossil find

In a diversification model with fossils, three types of event may occur on a lineage *ℓ* at a time *t*, namely:

1. a speciation: *ℓ* gives rise to a new lineage;
2. an extinction: *ℓ* no longer exists after *t*;
3. a fossil find: a fossil from *ℓ* is found (at contemporary time) and correctly dated at *t*, the actual time of burial of one of its specimens in the sediments.

The simplest way to model diversification and fossil find is to assume that the speciation, extinction and fossil find events occur at constant rates λ, *μ* and *γ* respectively. This model was first introduced and studied in Stadler (2010), which, among other results, provided the likelihood of the observable part of its realizations. In our recent study (Didier *et al*., 2012), we unfortunately were unaware of Stadler (2010) paper and expressed the same likelihood in a slightly different form, on which we will rely below. The model studied in Stadler (2010) is actually more general since the extant taxa are sampled with a probability *ρ*. For the sake of simplicity, we assume that we known all the extant species (i.e *ρ* = 1). Appendix ?? shows how to extend the computations to the case where extant taxa are represented by a non-exhaustive sample.

Formally, by identifying the state of the process at time *t* by the number *Z* (*t*) of lineages alive at *t* and *Y*(*t*), the cumulated number of fossil finds from the beginning of the diversification process to *t*, the joint process (*Z*(*t*), *Y*(*t*)) is defined as in Stadler (2010) and Didier *et al*. (2012)

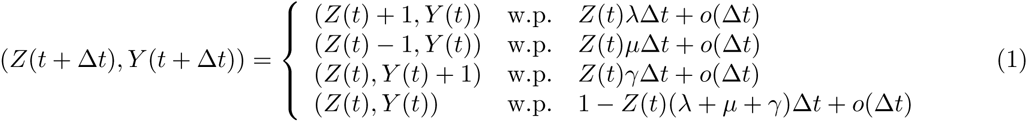

We set (*Z*(0), *Y*(0)) = (1,0) for the initial condition (i.e. the process starts from a single lineage and no fossil find).

## 3 Likelihood of reconstructed trees with fossils

### 3.1 Patterns

Previous works show how to compute the likelihood of a reconstructed tree with fossils with both divergence and fossil times (Stadler, 2010). Fossil dating relies on direct and physical evidence. But divergence times are always indirect measures, generally evaluated from molecular data that are available for extant taxa and Late Pleistocene fossils only.

Computing the likelihood of a reconstructed tree which is only known from its shape and from its fossil times, is thus an interesting question. We propose here a computational approach in which the likelihood of a reconstructed tree is decomposed as a sum-product of likelihoods of small parts of the diversification process, called *patterns.* Namely, a *pattern* 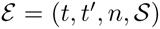 is a part of the diversification process with fossils which starts from a single lineage at time *t* and ends with *n* lineages and a tree shape 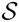 at time *t*′ (Figure 1). We shall consider only special classes of patterns which contain either no fossil or a unique one at their end time. They will be formally defined below and referred to as patterns of types a, b and c (Figure 1).

**Figure 1:**
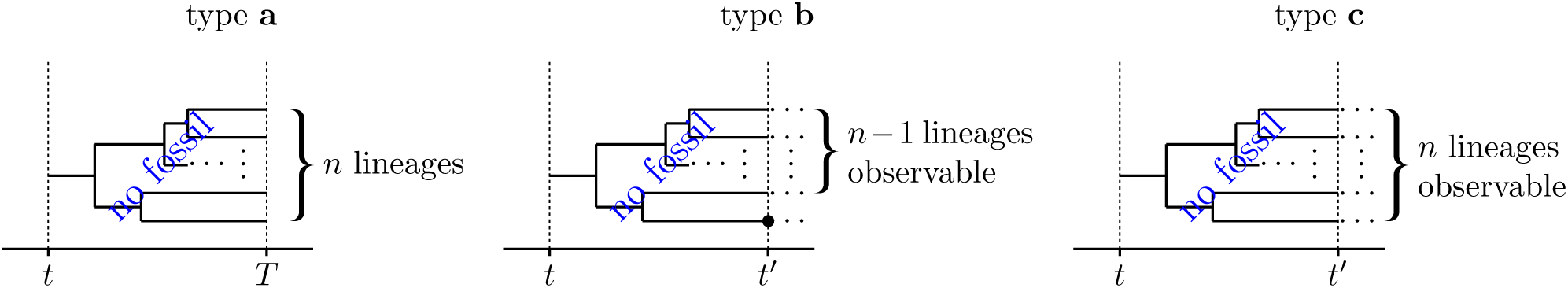
The three patterns on which our computations are based. The two trees to the right continue after *t*′.

### 3.2 Likelihood of ending with *n* lineages without observing fossils – Type a

#### Definition 1

*A pattern* 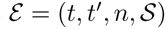 *of type* a *starts with a single lineage at time t, contains no fossil find and ends with n lineages and a tree shape* 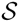 *at the contemporary time, i.e. t*′ = *T (Figure 1-left).*

In order to compute the likelihood of a pattern of type a, and for more general use, we shall compute the probability **P**(*n*, *t*) to have *n* lineages at time *t* without finding any fossil dated from 0 to *t* and by starting from a single lineage at 0, i.e. **P**(*n*, *t*) = ℙ((*Z*(*t*), *Y*(*t*)) = (*n*, 0)). With the definition of diversification process (1), **P**(*n*, *t*) satisfies the differential equations

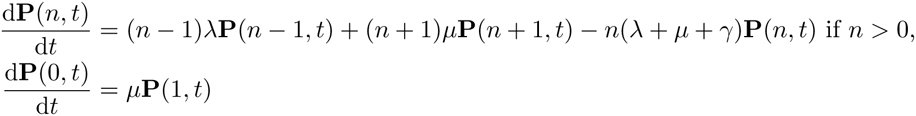

In Didier *et al*. (2012), we studied the generating function 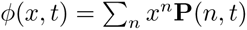 which satisfies

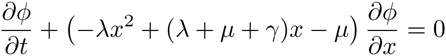

It was solved by using the method of characteristics to obtain

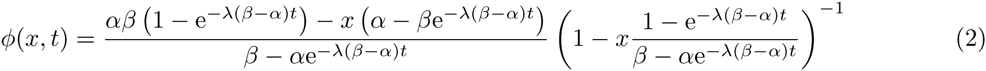

where *α* < *β* are the roots of −*λx*^2^ + (*λ* + *μ* + *γ*)*x* − *μ* = 0 which are always real (if *λ* is positive) and are equal to

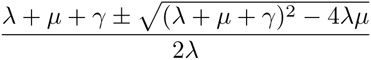

This quadratic equation comes in an other context and is further discussed in Appendix ??.

Let us set *ω* = − *λ*(*β* − *α*). Equation (2) may be equivalently written as

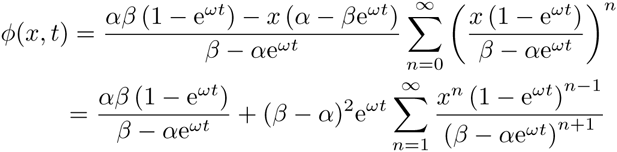

from which we finally get the probabilities we are looking for

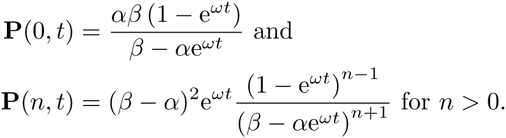

In particular, the probability for a lineage alive at time *t* to have exactly *n* ≥ 0 descendants at contemporary time without leaving any fossil between *t* and *T* is **P**(*n*, *T* − *t*) (Figure 1 Left).

#### Claim 1

*Under the diversification model with fossils, the probability of a pattern* 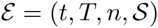 *of type* a *is*

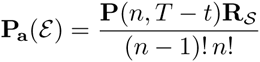

##### Proof

Since the diversification process is lineage-homogeneous and only births occur in a pattern of type a, the assumptions of Theorem ?? are granted (Appendix ??). It gives us the probability of the tree shape 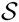 conditioned on having *n* leaves. The probability of 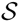 to have *n* leaves is that of 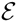 to end with *n* lineages, i.e. **P**(*n*, *T* – *t*).

### 3.3 Likelihood of an observable tree at its first fossil find – Type b

#### Definition 2

*A pattern* 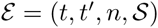 *of type* **b** *starts from a single lineage at t, ends with n lineages and a tree shape* 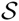 *at t*′ *and contains a unique fossil find dated at t*′ *(Figure 1-center).*

The probability **P**_o_(*t*) for a lineage to be observable at *t* is the complementary probability of being extinct before the contemporary time without any fossil find dated after *t*. We have (Stadler, 2010; Didier *et al*., 2012):

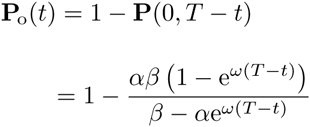

#### Claim 2

*Under the diversification model with fossils, the probability density of observing a fossil find at time t with n lineages alive at t and without any fossil find between* 0 *and t, by starting with a single lineage at time* 0 *is*

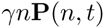

##### Proof.

This probability density is the product of the three following terms

- the probability to get *n* lineages at *t* without fossil find between 0 and *t*, which is **P**(*n*, *t*),
- the probability density to wait exactly 0 from *t* until the next event, which is *n*(λ + *μ* + *γ*),
- the relative probability of this event to be a fossil find, which is 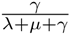.

Let now define **P**_x_(*k*, *t*, *t*′) as the likelihood of observing a fossil find at time *t*′ on a lineage in addition to *k* ≥ 0 other lineages observable at time *t*′, by starting from a single lineage at *t* (Figure 1-center). This likelihood may be obtained by summing over all numbers *j* ≥ 0, the likelihoods of observing no fossil find between *t* and *t*′, and *j* + *k* +1 lineages at *t*′ among which

1. a unique lineage has a fossil find dated at *t*′,
2. *k* lineages are observable at *t*′, not including the one with the fossil,
3. *j* lineages are not observable at *t*′.

We have

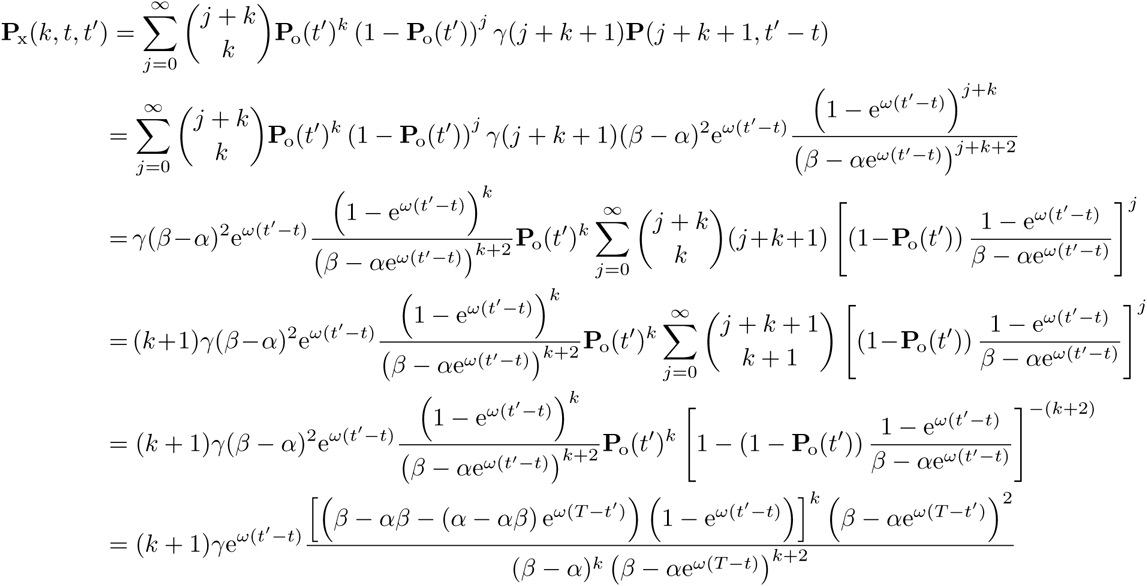

The likelihood **P**_x_(*k*, *t*, *t*′) is that of observing the ending configuration of a pattern 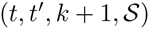 of type b. Since we are still dealing with pure-birth realizations of lineage-homogeneous processes, Theorem ?? gives us the conditional probability of a tree shape given this ending configuration.

#### Claim 3

*Under the diversification model with fossils, the likelihood of a pattern* 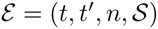 *of type* b *is*

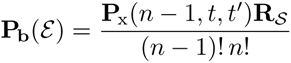

### 3.4 Likelihood of a reconstructed trees without divergence times

Patterns of type a and b will be the bases of our computations. Sometimes, we will consider their likelihoods conditioned on the fact that their starting lineages are observable at the times *t* when they begin. They will be referred to as likelihoods a^*^ and b^*^, respectively:

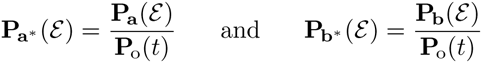

#### Theorem 1

*Under the diversification model with fossils, the likelihood of a reconstructed tree with fossil finds but without divergence times may be expressed as a sum-product of likelihoods* **a**, **b**, **a**^*^ *and* **b**^*^.

##### Proof

From the definition of the model, what is happening after a fossil find is independent of what was before it. In other words, the very same process as the one operating at the origin (re-)starts after each fossil find. Since they are all independent, the total likelihood of a realization may be expressed as a product of the likelihoods of the sub-parts of the realization obtained by splitting it at each fossil find (Figure 2). In this way, we obtain sub-trees, which will be referred to as *basic trees,* starting either at the process’ origin or at the fossil finds’ times and ending either at the contemporary time or a fossil find date. Note that, by construction, there is no internal fossil find in a basic tree. The starting lineages of the basic trees obtained from such splitting are not necessarily observable (these last ones may be empty or not; see Figure 2). In this case, we will talk about *unconditional basic trees.* Below, we will deal with another type of splitting leading to basic trees starting with observable lineages and which will be referred to as *conditional basic trees*.

**Figure 2.**
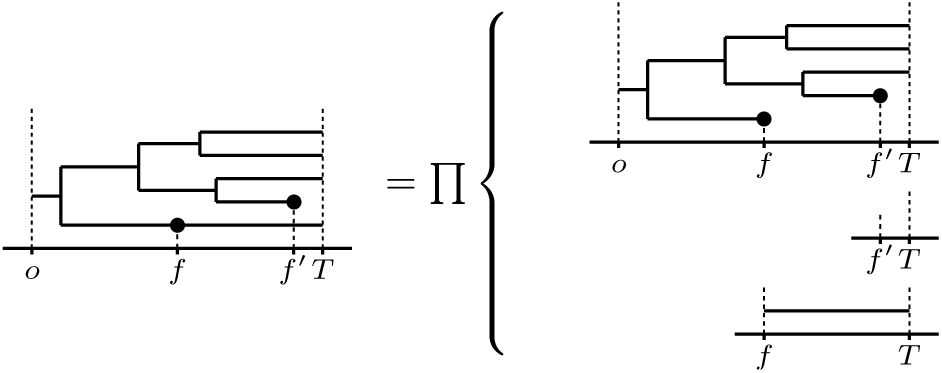
First stage of the likelihood computation decomposition of the reconstructed tree into trees with no internal fossil finds. The likelihood of the tree to the left is the product of that of the 3 trees to the right. The tree at the top right is the initial one cut at the first fossil finds encountered while the two others are those starting from each fossil find: the tree to the right center is a non-observable tree (i.e. the lineage is extinct with no fossil after *f*′) starting from a single lineage at time *f*′, and the tree to the bottom right is just a lineage starting from *f* to end at *T* without any observable event.

In order to prove that the likelihood of a basic tree may be expressed as a sum-product of likelihoods of types a, b, a^*^ and b^*^, we proceed by induction on the number of fossil finds contained in the basic tree. The base cases of our induction are of two types, namely:

1. the basic trees without fossils, which include the empty tree,
2. the basic trees made of only one lineage ending with a fossil find.

Since by construction, a lineage of a basic tree ends either at a fossil find or at contemporary time, trees of the base case 1 are necessarily patterns of type a, thus have likelihoods a or a^*^, depending on whether they are conditioned. The basic trees of case 2 are particular patterns of type b, thus have likelihoods b or b^*^, still depending on whether they are conditioned.

Let now assume that the theorem holds for all basic trees containing up to *m* fossils. Let 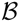 be a basic tree with *m* + 1 fossils and which is not among the base cases. In particular, it contains at least one fossil find. Let us refer to the time of the oldest one, which is almost surely unique, as the *first fossil time.* Since the tree is basic, this fossil find is on a leaf which is thus associated to the oldest known time of the tree but that of the origin. In order to compute the likelihood of 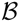, we are first interested in the relative positions of its nodes with regard to its first fossil time leaf. Let us consider all the possible assignments “before or after the first fossil time” of the nodes of 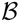. Namely, such an assignment is *possible* (i.e. consistent with the topology of 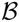) if:

- all the ancestor nodes of the leaf bearing the oldest fossil find are before the first fossil time (in particular the root of a basic tree is always before its first fossil time);
- all the leaves but that bearing the oldest fossil find are (strictly) after the first fossil time;
- if a node is before the first fossil time, then so are all its ancestors;
- if a node is after the first fossil time, then so are all its descendants.

Since the tree is finite, there are a finite number of such assignments. Let us next remark that any two before/after assignments are mutually exclusive. It follows that the likelihood of a basic tree is equal to the sum of the joint likelihoods with all its possible before/after assignments. This part of the calculus is illustrated in the two first columns of Figure 3. The first contains a basic tree and the second, all its possible before/after assignments with regard to its first fossil time.

**Figure 3:**
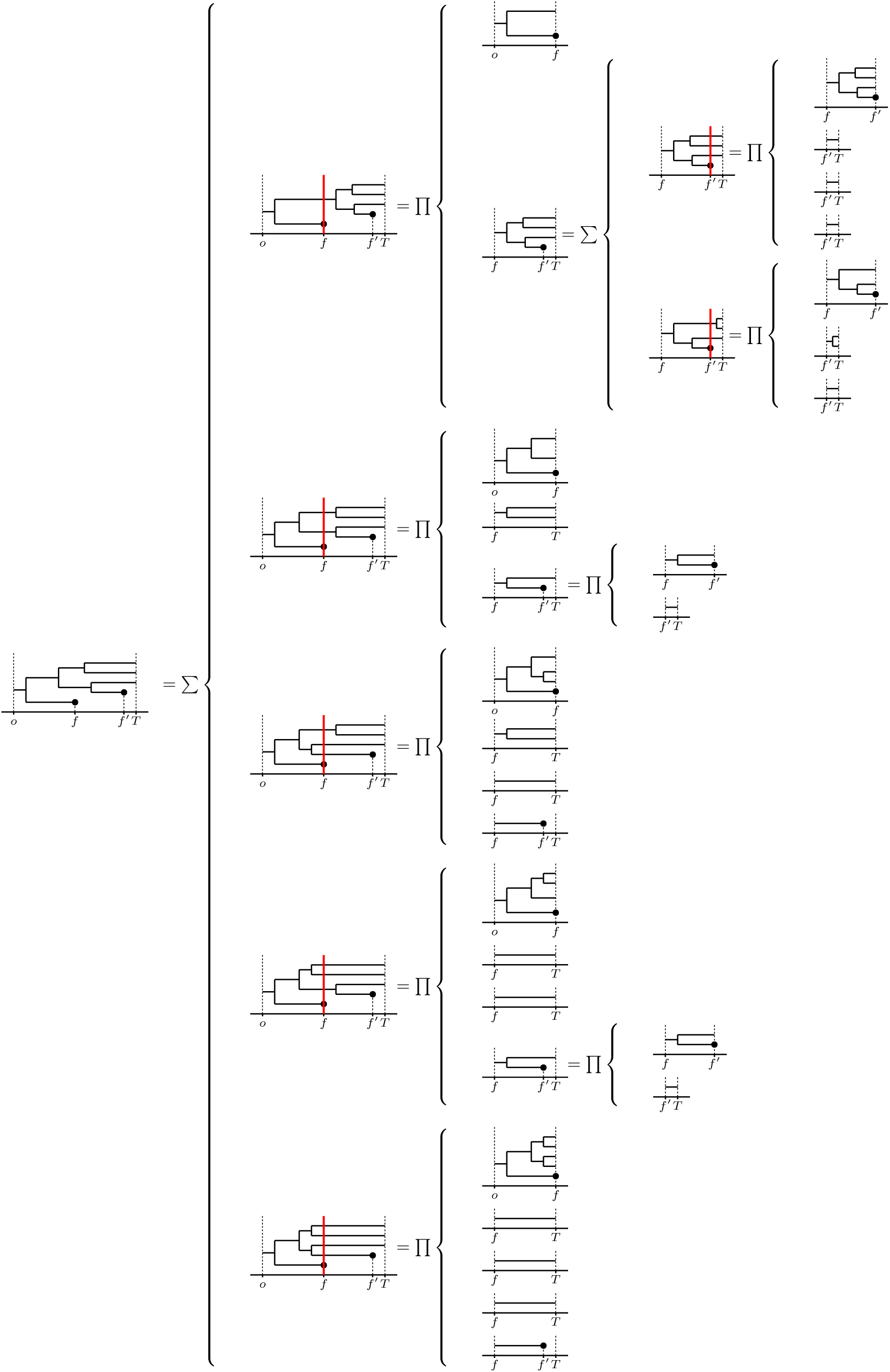
Stage 2 of the likelihood computation: the likelihood of the basic tree figured at the left-most is a sum-product of all the likelihoods of the right-most basic trees. Among these last ones, those starting from *o* are of types b and the remaining ones are of types b^*^ or a^*^ depending on whether they contain a fossil find. For all the trees containing fossil finds, the first fossil time is figured as a red line.

As one can see on column 2 of Figure 3, a before/after assignment naturally cuts internal branches of a basic tree according to the relative positions of their nodes with regard to the first fossil time. This splits the basic tree 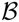 into:

1. the part of tree which is before the first fossil time (included), which contains the root and, by construction, has likelihood b or b^*^, depending on whether 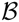 is conditioned;
2. what remains, namely a set of basic trees starting from the first fossil time with observable lineages (i.e. all these basic trees are conditioned).

Under the diversification model, what happens after the first fossil time, given the lineages observable at this time, is independent of what happens before. Moreover, all the basic trees starting from the first fossil time are independent one from another. The joint likelihood of the basic tree and one of its before/after assignment is thus the product of a likelihood of type b or b^*^ for the part before the first fossil time and of the likelihoods of the conditional basic trees starting from this time. This point is illustrated with the second and the third columns of Figure 3. For each tree/assignment of column 2, the parts resulting from its first fossil time splitting are displayed at the right of its brace. Among them, the top one is the part which is before the first fossil time, the others are for the conditional basic trees starting at the first fossil time of which likelihood computation is sketched in the next columns. Since they do not contain the oldest fossil of 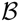, the basic trees starting from the first fossil time contain less than *m* fossil finds. With the induction assumption, their likelihoods may be expressed as a sum-product of likelihoods of types a, b, a^*^ and b^*^. The same thus holds for the likelihood of 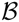 and, finally, of the whole tree.

The induction of the proof of Theorem 1 suggests a recursive computation of the likelihood, sketched in Algorithm 1. The complexity of the algorithm heavily depends on the tree shape and on the fossil density. The software *Diversification* provides a complexity index of a tree, which somehow predicts the time of its likelihood computation (Appendix ?? or the help of the software). Worst cases are balanced trees with few fossils for which the complexity is exponential with the size of the basic trees.

~~~
Function 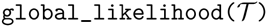
     Split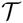 at each fossil find
     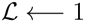
     forall the *split unconditional basic trees* 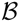 do
       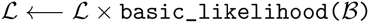
     return 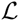
Function 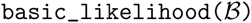
     if 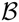 is a base case then
       return its likelihood
     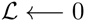
     forall the before/after assignments 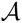 of 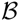 with regard to its first fossil time do
          Split 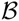 according to 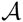
          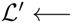 likelihood of the part before the first fossil time
          forall the split conditional basic trees 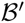 starting at the first fossil time do
            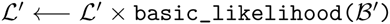
          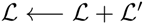
     return 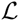
~~~

#### Algorithm 1

Computation of the likelihood of a reconstructed tree with fossil dates but without the divergence times. The first function returns this likelihood by calling the second one which recursively computes the likelihood of basic trees.

### 3.5 Dealing with missing data – Type c

It may occur that one knowns for sure that a lineage is alive (and observable) at a given time without having enough information to determine the part of the phylogenetic tree descending from it after this time, for instance because of the quality of the fossil record, or because a database was compiled within a given time interval ending in the past (not reaching the present). We will say that the lineage is *not considered after* this time, which will be referred to as a *stopping time* of the tree. A noticeable difference between fossil and stopping times is that, under the model, dating two fossils with a same time has a null probability, whereas there is no objection to set several stopping times at a same date (indeed, this is likely if a database was compiled between the origin of a clade and the end of a given geological stage).

Basically, stopping times are not events of the diversification process with fossils, but rather inform about the limits of the database that is available to estimate parameters.

The likelihood of the reconstruction of a tree with such missing data is computed in a very similar way as above. It needs to consider an extra type of patterns.

#### Definition 3

*A pattern* 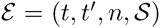 *of type* **c starts with a single lineage at** *t and ends with n* > 0 *lineages observable at time t*′ *and with the tree shape* 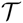 *without observing any fossil find meanwhile (Figure 1-right).*

Let **P**_y_ (*k*, *t*, *t*′) be the likelihood of ending with *k* > 0 lineages observable at time *t*′ by starting from a single lineage at *t* without fossils meanwhile. We have

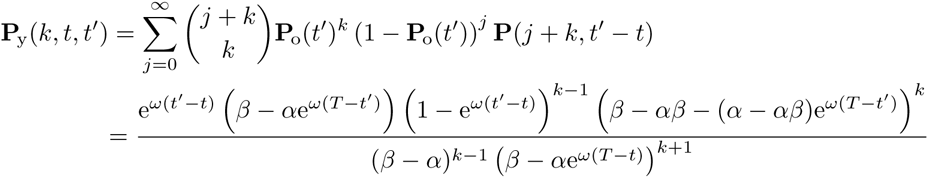

In the same way as with Claims 1 and 3, **P**_y_(*k*, *t*, *t*′) is the likelihood of the ending configuration of a pattern 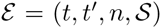 of type **c**, which has to be multiplied by the conditional probability of the tree shape 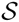 to get the likelihood of 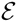. Patterns of type **c** satisfy the assumptions of Theorem ??, which gives us the following claim.

#### Claim 4

*The likelihood of a pattern* 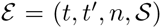 *of type* **c** *is*

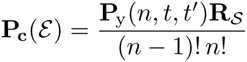

We put 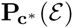 for the conditional likelihood of E given that its starting lineage is observable at *t*.

**Theorem 2.** *Under the diversification model with fossils, the likelihood of a reconstructed tree with fossil finds but without divergence times containing missing data may be expressed as a sum-product of likelihoods of types* **a**, **b**, **c**, **a**^*^, **b**^*^ *and* **c**^*^.

##### Proof.

The proof follows the same outline as that of Theorem 1. By splitting a reconstructed tree with both fossils and missing data, at each fossil find, we get basic trees which still starts from the origin or fossil dates but now ends either at the contemporary time, at fossil dates or (that is a new possibility) at stopping times.

In order to prove that the likelihood of a basic tree has the desired form, we proceed by induction on the number of fossil and of stopping times. The base cases are now

1. the basic trees which contain neither fossils nor missing data (and possibly no lineages at all),
2. the basic trees made of only one lineage ending with a fossil find,
3. the basic trees only made of lineages with an unknown fate after a same stopping time.

Already considered with Theorem 1, the first two cases lead to likelihoods a, a^*^, b or b^*^. The basic trees of the third case fit the definition of patterns of type c, and thus have likelihoods c or c^*^, depending on whether they are conditioned.

Let us consider a basic tree 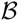 starting at *t*, which is not a base case. It contains at least a leaf which is not contemporary. Let us assume that its oldest leaf time *t*′ is a stopping time (otherwise the situation is handled as in the proof of Theorem 1). A before/after assignment of 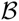 with regard to *t*′ splits it into

- the part before *t*′ which is a pattern of type c, thus has a likelihood c or c^*^ depending on whether 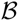 is conditioned,
- the part after *t*′, which is the set of the basic trees starting at *t*′ from all the lineages observable at this time except the ones that are not considered after *t*′ (there is at least one of them but there may be possibly more).

The joint likelihood of 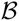 with a before/after assignment is thus a product of likelihood of type c or c^*^ and of likelihoods of basic trees which contain at least one stopping time less than 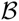. Since the likelihood of 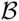 is a sum of such products, the induction assumption allows us to conclude that it has the desired form.

Modifying Algorithm 1 in order to deal with missing data is straightforward.

## 4 Diversification rates estimation - Simulation results

In order to assess the estimation approach we follows the same protocol as in Didier *et al*. (2012). We first simulate artificial evolution under the diversification model with fossils with given speciation, extinction and fossilization rates. Let us first remark that several sets of data, varying in information content and kind, can be taking into account from a simulated evolution:

A. the “complete information” that is the whole tree, fossils and all event times, including timing data about extinct lineages with no fossil finds (Figure 4-A),
B. the phylogeny from the reconstructed tree with fossils including the divergence times of both extant and extinct taxa and the fossil ages (Figure 4-B),
C. the phylogeny from the reconstructed tree with fossils including only the fossil ages as temporal information (Figure 4-C),
D. the phylogeny of the extant taxa, with their divergence times (Figure 4-D).

**Figure 4:**
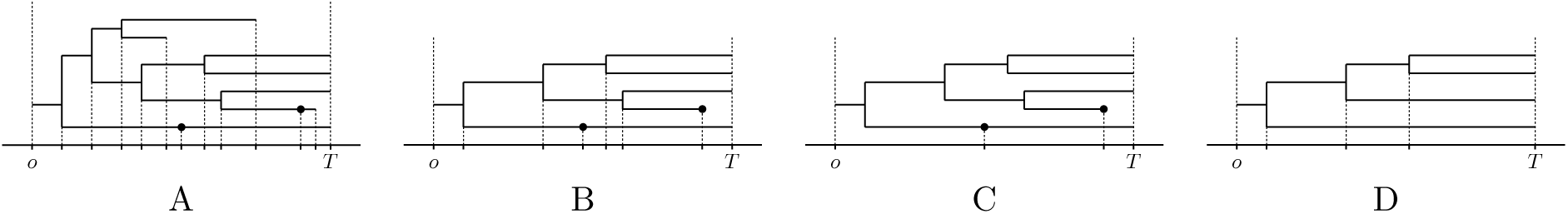
The various kinds of information from which we estimate the diversification rates. The trees are sorted from those for which we use the most to those for which we use the least data, except for the last two, which use two non-overlapping temporal information. This order of tree representations also corresponds with those that yield the least (left) to those that yield the greatest (right) error in parameter estimates.

Under the diversification model with fossils, the likelihood of trees with these various sets of time data has been derived. The likelihood of the complete information (A) is obtained by counting the number of each type of event (Keiding, 1975). The likelihood of the reconstructed tree with fossils (B) was derived in Stadler (2010). The same one but without the divergence times (C) is the main object of this work. Last, the likelihood of the phylogeny of extant taxa (D) was provided in Nee *et al*. (1994).

We are thus able to perform maximum likelihood estimation of the rates from the various kinds of datasets described above. Since the set D does not take into account fossil finds at all, the fossil rate is estimated only for sets A to C.

In practice, we simulate trees with a fossil record in the same way as in Didier *et al*. (2012). For technical reasons, we filter the simulated data obtained according several criteria. We reject all the trees ending with less than 20 extant taxa or with more than 1000 nodes. In order to limit the total computation time of our evaluation protocol, we keep only simulated trees with a complexity index under a fixed threshold set here to 5.10^7^ (Appendix ??). The maximum likelihood estimation is performed on the filtered simulated data by using the numerical optimization algorithm BOBYQA (Powell, 2009) of the *NLopt* library (Johnson, 2015).

Figure 5 displays plots of the mean absolute error in the maximum likelihood estimation of the speciation, extinction and fossil find rates for the various kinds of datasets extracted from the same set of simulated trees. We observe that, as expected, estimating from fossil ages only leads to a greater mean absolute error than estimating from the whole information available or by taking also into account the divergence times, but these estimates are still fairly accurate. A very interesting point here is that, with the fossil rates used to simulate data, using only fossil ages (C) always leads to more accurate estimates than using the phylogeny of extant taxa with divergence times (D).

**Figure 5:**
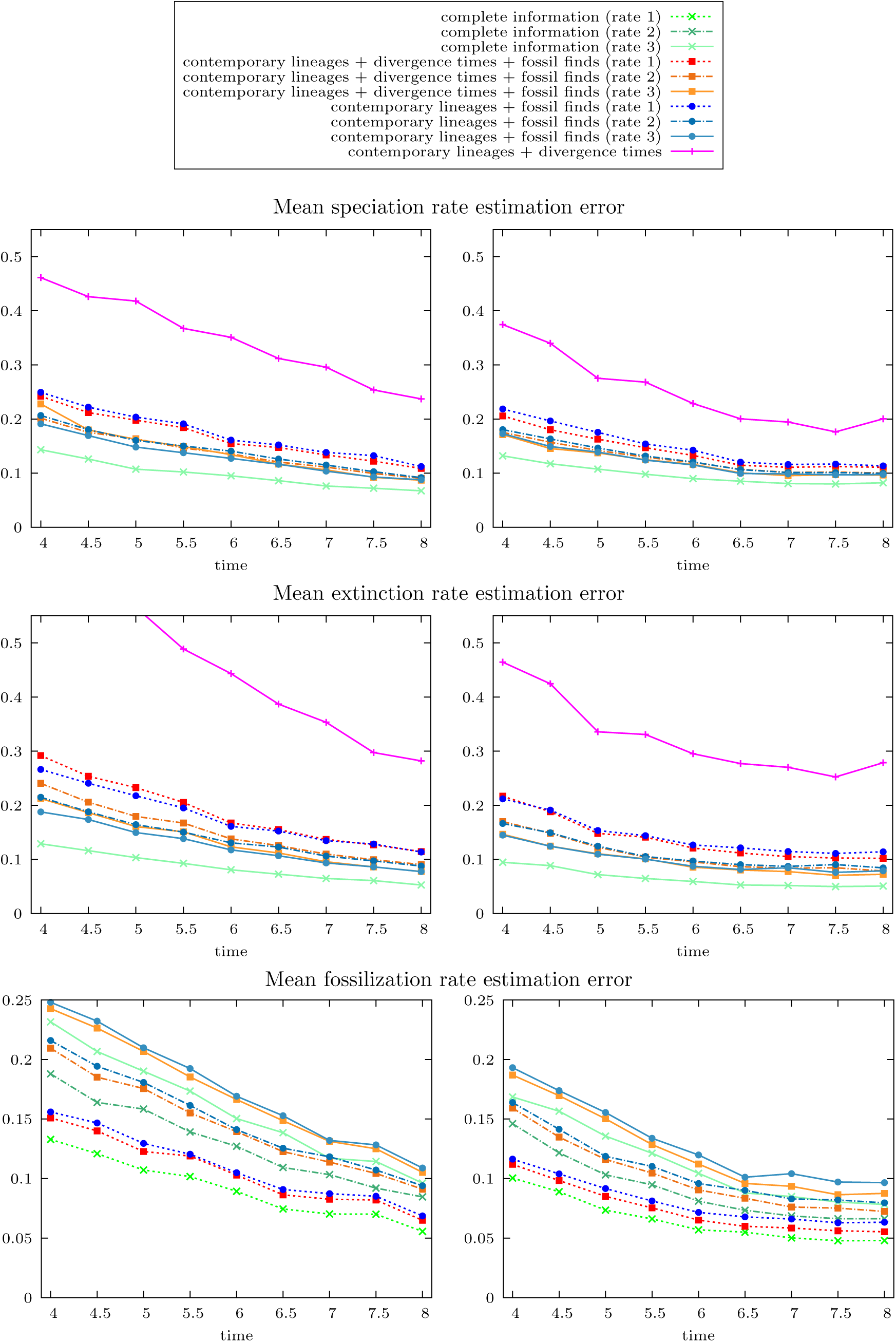
Mean absolute error of speciation (cladogenesis), extinction and fossil discovery rate estimation *versus* the simulation time over 1000 simulated trees with 1.5/1 (Left) and 2.0/1 (Right) speciation/extinction rates and fossil rates from 1 to 3.

## 5 Empirical example

### 5.1 Dataset compilation

We provide an empirical example with a dataset of Eupelycosauria, a clade that includes mammals and most of their stem-group (Reisz, 1986); the only synapsids (the total group of mammals) excluded from eupelycosaurs are the Caseasauria (Figure 6). We used the data matrix from Benson (2012) (because this is the matrix with the greatest number of relevant terminal taxa that we know of) and added the taxa that were not included in the matrix into the tree based on the literature, which is reviewed in Benson (2012). This tree, provided in SOM (Supplementary On-line Material) 1 is only one out of 200 000 most parsimonious trees (there were more, but this was the limit we entered, to avoid a crash), so it should not be viewed as more than one of many plausible trees for this clade. Indeed, Benson (2012) also obtained a poorly resolved strict consensus tree with his complete dataset, and could only increase its resolution by pruning taxa. But our analysis, on the contrary, required adding all reasonably well-known taxa. The detailed stratigraphic data are provided in SOM 2, and the references in SOM 3.

**Figure 6:**
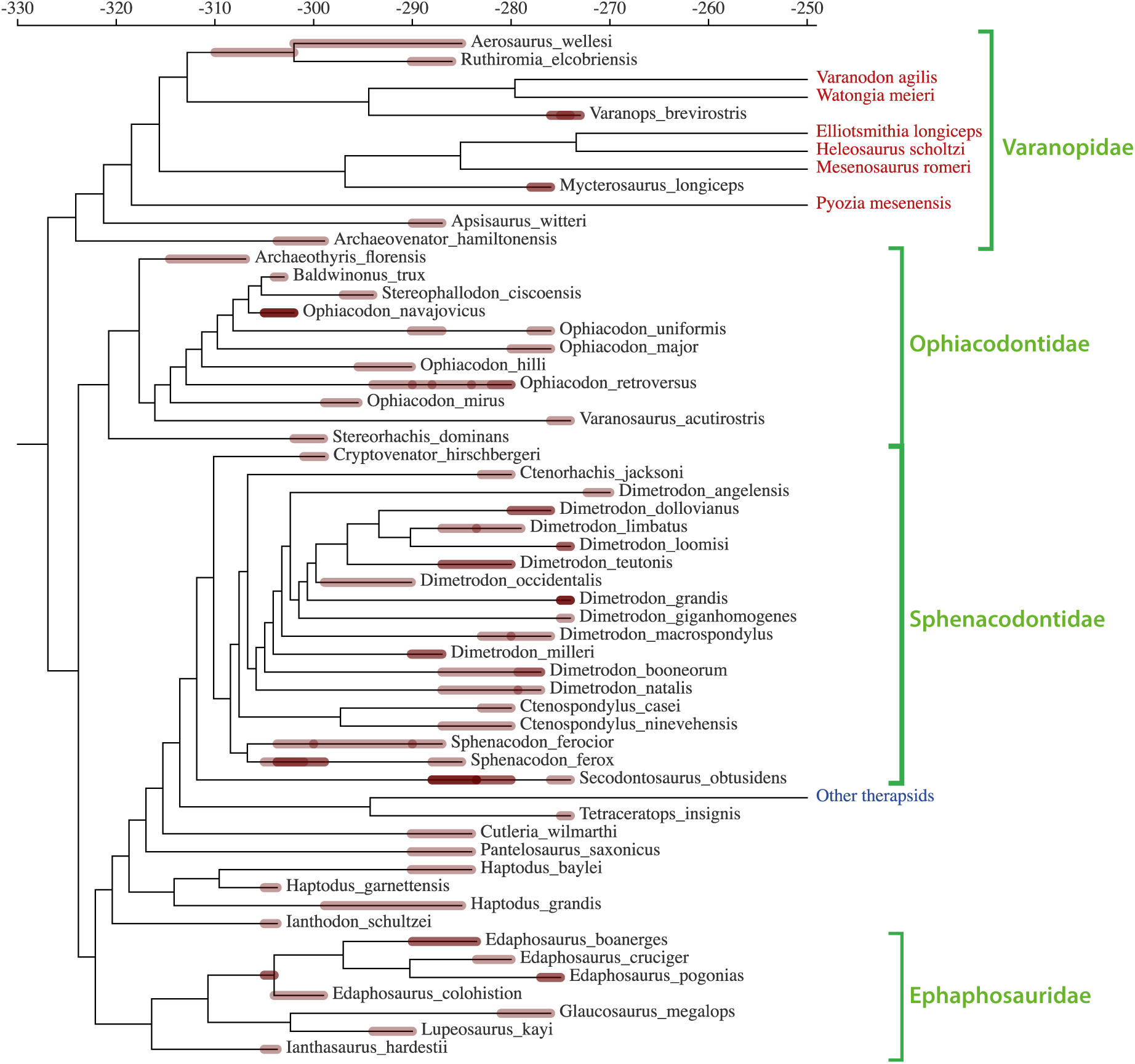
Tree of Eupelycosauria. The taxon “Other therapsids” is extant; the other taxa lacking a fossil record before 270 Ma have an unknown fate after 270 Ma (most recent time for which fossil data were collected into our database). Two taxa considered ancestral here (for illustration purposes) appear as fossil occurrences on internal branches. These are *Edaphosaurus novomexicanus* and *Aerosaurus greenleorum*.

Our model, and indeed previous analyses (Foote, 1996), predict that some actual ancestors should be found in the fossil record, but parsimony analyses do not identify them, since the resulting cladogram has all taxa on terminal branches. Thus, we used MacClade’s (Maddison and Maddison, 1992) “Make ancestor” command to verify if some taxa could plausibly be considered ancestors. For this, three conditions had to be met: first, the ancestor needs to be older than its presumed descendants; second, making the taxon an ancestor could not increase the number of parsimony steps; third, presumed ancestor and at least one of its descendants had to be scored into the matrix (rather than having their systematic position having been determined from the literature). Only two taxa passed this rather stringent test: *Edaphosaurus novomexicanus,* which may be ancestral to four congeneric species (*E. colohistion, E. boanerges, E. cruciger,* and *E. pogonias*), and *Aerosaurus greenleorum,* which may be ancestral to *Aerosaurus wellesi* and *Ruthiromia elcobriensis.* However, note that this is better-supported for *E. novomexicanus,* which is scored for 113 phenotypic characters, than for *A. greenleorum,* which is scored for only 26 characters. These two ancestral taxa are not identified on the tree but their presence is indicated by the fossil occurrences on the relevant internal branches (Figure 6).

### 5.2 Estimating diversification rates from empirical data

Dating fossils is quite a complex process. In practice, the geological age of an individual fossil is given as an interval of time determined according to the geological layer where it has been found, and which has typically been indirectly dated through stratigraphic correlations with other strata that bear radioisotopes, or through magnetostratigraphy (Gradstein *et al*., 2004). In the same way, the age of the beginning of the diversion is never precisely known and consequently, in our software, it may be given as a range of possible ages.

Nevertheless, our estimation of diversification rates is based on likelihoods which are computed with exact times. In order to apply our approach in a practical situation, we turn each time interval of the empirical data into an exact time which is uniformly sampled from the interval of possible ages that has been specified for each fossil and the diversion origin. In order to assess the robustness of the estimates thus obtained, we repeat this sampling process several times (here, 1000) and consider the means and the standard deviations of the results.

### 5.3 Diversification rates of eupelycosaurs

Our software estimates the speciation (cladogenetic) rate at 1.005E-1, with a SD (Standard deviation) of 2.954E-3, an extinction rate of 9.622E-2 with a SD of 2.828E-3, and a fossilization rate of 3.122E-2, with a SD of 9.174E-4. All rates are in events/My, when we consider that *Edaphosaurus novomexicanus* and *Aerosaurus greenleorum* are ancestors and that the divergences between taxa that occur after the studied interval (in red in Figure 6) had occurred within the interval (before 270 Ma). However, considering that these divergences (whose timing is poorly constrained) occurred after the studied interval changes only marginally the results: speciation rate of 9.594E-2 with SD of 2.895E-3, extinction rate of 9.488E-02 (SD of 2.863E-3), and fossilization rate of 3.214E-2 (SD of 9.697E-04). Even the log-likelihoods are similar in both cases, at -553 in both cases. This case can be assessed by leaving only one branch per clade that occurred after the studied interval. For instance, in the clade that includes *Varanodon agilis* and *Watongia meieri*, one of these two taxa is pruned. Similarly, considering that *Edaphosaurus novomexicanus* and *Aerosaurus greenleorum* are not ancestors changes the results only marginally: speciation rate of 1.090E-1 with SD of 3.173E-3, extinction rate of 1.047E-1 (SD of 3.048E-3), and fossilization rate of 2.949E-2 (SD of 8.579E-4), with a log-likelihood of -595.

Using the formulae in Appendix D, we determined that the probability that a lineage in the clade of interest and in the studied timeframe left no observable fossil, either on its base branch, or on all its descendants, is 0.5608. This translates in a probability of only 0.1370 that a given branch (still defined as an internode in a tree) of the real tree (only part of which is visible) is directly documented in the fossil record by at least one fossil (in both cases, we have rounded off the numbers to the fourth decimal).

## 6 Discussion

Our most interesting and surprising find is that we can estimate speciation and extinction rates more accurately using the age of fossils found than by using divergence dates. This result raises the possibility that further development of our method could lead to a much better understanding of the evolution of biodiversity through time because uncertainly about the age of fossils is much less, typically, than uncertainty about divergence dates. For instance, Hugall *et al*. (2007) report 95 percent confidence intervals of 304-342 Ma and 263-321 Ma around the best estimates of the ages of Lissamphibia (323 or 292 Ma, depending on data used), which is reasonably precise, but some much younger clades, like Pleurodira (best estimates from 114 to 182 Ma, depending on source data) had proportionally wider 95 percent confidence intervals (92-136 Ma and 144-220 Ma, respectively). Similarly, in a more recent, total evidence (tip) dating study, Pyron (2011) reported a 95 percent credibility interval of only about 27 My for Lissamphibia (which he dated to about 300 Ma), which is again fairly precise, but other nodes were estimated much less precisely, such as Gymnophiona (dated at 75 or 98 Ma, depending on the analysis, with 95 percent credibility intervals of 40-110 Ma and 19-206, respectively) or Anura (198 or 226 Ma, with intervals of 157-238 and 159-276 Ma, respectively). These much wider credibility intervals presumably better reflect phylogenetic uncertainty of the fossils used to calibrate the tree, as well as uncertainty about the length of branches subtending extinct taxa. This is coherent with similarly wide credibility intervals reported by Ronquist *et al*. (2012b) for Hymenoptera, with a 95 percent credibility interval of the root age of 320-410 Ma. In that study, the node dated with the greatest relative precision (located just above the root) has a 95 percent credibility interval of 300-355 Ma, which encompasses most of the Carboniferous (299-359 Ma; Gradstein *et al*., 2012).

The geological age of fossils is generally much more precisely known, typically at the geological stage level, with many other, more problematic cases in which the range of plausible ages encompasses two consecutive geological stages. There are several geological stages per period; for instance, the Carboniferous (a period) encompasses seven geological stages (also called ages), from Trounaisian to Gzhelian (Gradstein *et al*., 2012). Thus, the age of the oldest known amniote, which is often used to calibrate molecular trees, is about 315 Ma, add or take 5 My (Davies *et al*., 2006). The exact width of the 95 percent confidence interval is difficult to assess because most of the uncertainty is attributable to uncertain correlations with the strata that were dated through radiometric methods, as well as to interpolation between radiometric dates, which may be separated from each other (at least in the Paleozoic) by a few million years. In any case, the strata containing the oldest known amniote fossils (in the Joggins Formation from Nova Scotia, Canada) were dated from the late Langsettian (Westphalian A) equivalent, late Bashkirian (Hower *et al*., 2000) through stratigraphic correlations using pollens with other strata where radiometric dating was performed, but all the dates given in the primary paleontological literature in the last three decades vary between 310 and 320 Ma, which encompasses part of two consecutive geological stages (Moscovian and Bashkirian) of the Carboniferous (Gradstein *et al*., 2012). The current estimate is near 318, using the latest geological time scale (Richards, 2013).

The uncertainty about nodal ages in molecular dating, at least those obtained using node dating, as opposed to total evidence or tip dating, have typically been under-estimated because most molecular dating studies, except for a few recent ones, did not incoporate uncertainty about the phylogenetic position of the relevant fossils, and this can drastically increase uncertainty of node ages (Sterli *et al*., 2013), although this would also affect the accuracy of our method). The greater uncertainty about divergence dates than about age of fossils is a consequence of the fact that molecular dates are obtained through calibration of molecular branch lengths using divergence dates whose ages are typically estimated through the fossil record or from geological events. We can never be certain that we know the oldest organisms that belonged to a taxon; indeed, several methods were developed to estimate the width of the confidence interval of the true stratigraphic range of taxa (Strauss and Sadler, 1989; Marshall, 1994, 1997, 2008; Marjanovic and Laurin, 2008; Wilkinson and Tavaré, 2009), but we are still working on better methods to estimate these (Laurin, 2012; Stadler and Yang, 2013; Heath *et al*., 2014). Thus, divergence dates are inherently more uncertain that the ages of fossils. Geological events are even less reliable to date trees because it is often difficult to be certain that a divergence between taxa is causally linked to such events, whose age is also uncertain.

The recent method of tip dating (also known as total evidence dating) makes different assumptions (Pyron, 2011; Ronquist *et al*., 2012a,b), but does not necessarily reduce uncertainty about divergence dates because it requires estimating branch lengths leading to extinct taxa from phenotypic characters whose evolution must be modeled using methods similar to those developed for molecular data. The method has the advantage of not requiring to fixing a priori node ages based on the fossil record (an advantage, because fossils only yield the minimal age of nodes), but estimates of the length of branches leading from tips (representing extinct and extant taxa) to nodes necessarily involve substantial uncertainty. There are both practical and theoretical reasons for this. Among the most obvious practical ones, the morphological datasets are much smaller than the average molecular dataset because morphological data are time-consuming to collect; thus, stochastic effects on branch length estimates for extinct taxa should be greater than for extant taxa, for which molecular data are available. Among the theoretical problems is the fact that morphological characters are much more complex than single nucleotides or aminoacids, and they are not as readily partitioned. Partitioning schemes could be devised; for the vertebrate skeleton, cranial vs. postcranial, dermal vs. endochondral, or axial vs. appendicular skeletal components could be considered as tentative partitions, and similar partition schemes could probably be devised for most taxa or organs, but their usefulness in this context remains to be assessed. To sum up, molecular (node) and total evidence (tip) dating involve substantial uncertaintly. Consequently, the prospect of no longer requiring divergence dates (other than for the root, which our method still requires) to estimate speciation and extinction times should represent an important breakthrough in this field.

The main caveat of our system is that placing precisely fossils into a phylogeny is typically more difficult than for extant taxa because only phenotypic data (mostly morphological, sometimes histological) are accessible in fossils more than about 0.1 Ma. Furthermore, compiling large data matrices of phenotypic data is more time-consuming than for molecular data because there are no databases equivalent to GenBank (http://www.ncbi.nlm.nih.gov/genbank/), even though some databases of different kinds exist both for the fossil record, such as the Paleobiology database (https://paleobiodb.org/#/) and for phenotypic data, such as MorphoBank (http://www.morphobank.org/). However, paleontologists increasingly compile such data matrices to tackle a variety of related questions about the evolution of taxa, as shown by the recent inauguration (in 2003) of the Journal of Systematic Palaeontology (http://www.tandfonline.com/toc/tjsp20/current) and its subsequent success. Thus, our method will both enable the scientific community to tap into these increasingly-available phylogenetic data, and allow paleontologists to extract more evolutionary information out of their data matrices.

Our simulations assume that both temporal (divergence, fossil age) and topological (phylogenetic) information is correct (without error). Of course, this is never the case in empirical datasets. The fact that phylogenetic uncertainties about fossils are presumably greater than for extant taxa is compensated, at least in part, by the fact that the geological age of fossils is known with a greater precision than divergence times. It is difficult to determine how both factors would influence our results if they were taken into consideration, though they might more or less compensate each other. This topic would deserve being investigated, but is beyond the scope of our study.

The possibility of directly using fossil data to infer speciation and extinction rates should also allow, with further developments, to better study phenomena such as the timing of evolutionary radiations and mass extinction events. These have been studied recently with some methods that exploit molecular timetrees (Alfaro *et al*., 2009; Rabosky, 2014) or even asymmetry of uncalibrated ultrametric trees (Chan and Moore, 2005), but exploiting the fossil record should yield more reliable estimates, simply because much more data are potentially available through the fossil record, such as evolutionary radiations and mass extinction of completely extinct clades. The potential gain in precision should be especially great for extinction rates, which are notoriously difficult to estimate from extant taxa alone (Paradis, 2004). Much remains to be done in this field. So far, our method estimates only one speciation, extinction, and fossilization rate over a tree, but tackling the complex history of biodiversity changes through time will require detecting when shifts in these values occur in trees. If and when such methods are developed, they will also require new databases to be developed because currently-available compilations, such as those found in the Paleobiology Database (https://paleobiodb.org/#/), lack the detailed phylogenetic data required for the methods mentioned above.

In the meantime, we provide a simple application example (Eupelycosauria) that yields the first biological results from our method. These results are rather encouraging because the estimates have a small standard deviation, in the order of 3 percent of the estimated values, which suggests that they are fairly reliable. The estimated speciation rate (about 0.1/My) may seem low (implying a mean time between cladogeneses of about 10 My), but it is not necessarily wrong because two factors may have distorted our previous views on the speciation rates of Permo-Carboniferous synapsids. First, there may be too many nominal species; some of them may represent a chronospecies, a single lineage evolving through time (which is here considered a single species), while others may have been erected simply because they occurred in a different stratigraphic level or geographic area, or because of small morphological differences that may reflect ontogenetic stage rather than taxonomic status (Falconnet, 2015). Second, our estimates imply that only about fourteen percent of the Permo-Carboniferous eupelycosaur lineages (defined as an internode on the tree) have left a fossil record, so if our estimates of the parameters are more or less correct, the number of lineages that were diversifying at the time has probably been strongly under-estimated.

The possibility also exists that our parameter estimates are biased because the real rates of speciation or extinction may not have been homogeneous over time. Indeed, if the rates had been homogeneous, it would be surprising that of the lineages represented in our database, a single one (identified as “other therapsids” in Figure 6) extends to the present. The Early Permian is not generally considered to have witnessed mass extinction events, contrary to the Late Permian (Benton, 2003), but a somewhat less spectacular crisis appears to have taken place in the Middle Permian, around 260 Ma (Retallack *et al*., 2006; Wignall *et al*., 2009), shortly after the studied time interval. This event, or perhaps other as yet undetected still earlier and less dramatic events, may have caused significant heterogeneities in extinction rates that our model does not yet allow to account for. In any case, these mass extinction events help reconcile our parameter estimates with the fact that a single lineage persists today. Another possible confounding factor may be the appearance of the amniotic egg, which has long been seen as a key evolutionary event (Romer, 1957; Laurin, 2004), and which may have caused a rapid evolutionary radiation in the Carboniferous. The speciation rates may have decreased thereafter. A rapid evolutionary radiation followed by a decrease in diversification rate might create problems (whose severity we have not assessed) with the current implementation of our method, and will be addressed in future works.

The fact that we interpret (provisionally, pending further analyses, and for illustration purposes) two taxa (*Edaphosaurus novomexicanus* and *Aerosaurus greenleorum*) as plausible actual ancestors will surprise some paleontologists. There is a debate about whether or not ancestors (other than the trivial case of successive populations of a given lineage being preserved in several strata in the fossil record) can be identified in the fossil record. Until the 1960s, most paleontologists focused on searching for ancestors, and often claimed to have found them (Romer, 1966). This was often done prematurely, which led Hennig (1965) to protest energically against this practice. We concur, without rejecting the possibility of encountering actual ancestors in the fossil record. Our position is supported both by simulations (Foote, 1996) and by theoretical considerations (Bonde, 2001). Early studies of Permo-Carboniferous eupelycosaurs indeed suggested that some of the known taxa were probably ancestral to others; see, for instance, pages 215 and 230 in Romer and Price (1940). More recent studies on this set of taxa have been less optimistic about finding ancestors. Thus, Modesto and Reisz (1992) identified two autapomorphies in *Edaphosaurus novomexicanus* and concluded, on that basis, that it was not an ancestor of geologically more recent species of *Edaphosaurus* (contrary to the assumption we made on the illustrated tree). That may well be, but we note that reversals no doubt occur in evolution, so the presence of one or two autapomorphies does not necessarily indicate that a taxon is not ancestral; it only reduces the probability that it is. Using a model-based approach, it should be possible to estimate the likelihood of both cases (a taxon being sister-group or ancestral), but this would require substantial developments that are beyond the scope of our study. In any case, our results change only marginally when we consider that *Edaphosaurus novomexicanus* and *Aerosaurus greenleorum* are not ancestral, so in the present case, the ancestral (or not) status of these taxa is only marginally relevant, and this is fortunate because the identification of ancestors in the fossil record remains problematic.

## Author contribution

GD provided the general idea of the likelihood computation, derived the mathematical results, developed the algorithms and the software, ran the simulations and wrote much of the paper. ML contributed to the writing of the introduction, discussion, and presentation of the empirical example, supervised MF when she compiled the empirical dataset on eupelycosaurs, provided feedback about simulation settings and features to implement in the software, and edited the text for style, grammar and clarity.

## Acknowledgments

We thank Pauline Drapeau, who worked on an early version of the database from which our empirical example was extracted and reworked.

## Appendices

### A Probability of a tree shape

#### Number of birth rankings consistent with a tree

We are interested here in tree shapes (i.e. trees without time information) resulting from realizations of general birth and death processes. Most of the ideas of this section are close to those developed in (Ford *et al*., 2009).

To keep things as general as possible, we define a *realization* of n lineages with birth times *t*_1_ < *t*_2_ < … < *t_n_*_–1_ in the following way. The process starts with a single lineage at a time *t*_0_ < *t*_1_. At time *t_i_*, a lineage is picked among the lineages alive to give birth to a new lineage. Each lineage is associated to a different label in an arbitrary way (i.e. not depending on its birth date, its parenthood etc.). The natural (and usual) way to associate a tree shape with a realization is as follows:

- the internal nodes and the leaves of the tree are in one-to-one correspondence with the birth events and the lineages of the realization, respectively;
- the direct ancestor of the leaf associated with the lineage *x* is the node corresponding to the last event involving *x* (as parent or child);
- the direct ancestor of the internal node associated with the birth of the lineage *x* is the node corresponding to the last event before the birth of *x* that involves the parent of *x*.

By construction, the trees resulting from realizations are rooted, binary and (leaf) labeled (labels of leaves are those of the lineages).

The *scenario* of a realization is its sequence of birth events ordered following their occurrence times. The *i*^th^ event of a scenario *E* is noted *E_i_* and is of the form “lineage *x* is borne from lineage *y*”, *x* and *y* being referred to as the child and the parent lineages of *E_i_*, respectively. A given scenario is *valid* if there exists a realization from which it arises. Basically a scenario is valid if and only if all its lineages but the starting one are the child lineages of the earliest event involving them.

Several evolutionary scenarios may lead to a same tree. For instance, a tree made of only two lineages *x* and *y* may result from both the scenarios “*x* was borne from *y*” or “*y* was borne from *x*”. Conversely, the scenario of a realization fully determines its tree in the following way. If a tree shape 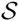 and a scenario *E* both result from a same realization, the internal nodes and the leaves of 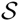 are in one-to-one correspondence with the events and the lineages of *E* respectively. The direct ancestor of the leaf associated with the lineage *x* is the internal node corresponding to the event with the greatest rank in *E* involving *x*, while the direct ancestor of the node associated with the event *E_i_* is the node corresponding to the event with the greatest rank strictly smaller than *i* which involves the parent lineage of *E_i_*.

Let **r** birth ranking of the lineages of a realization (i.e. **r** is the vector in which the *i*^th^ entry *r_i_* contains the *i*^th^ oldest lineage). A scenario *E* perfectly determines the birth ranking of its lineages: the starting lineage has rank 1 while the ranks of the other ones are obtained by adding 1 to the ranks of their birth events in *E*.

##### Remark A1.

*If a tree shape and a scenario come from a same realization, then the node n associated with the event “x is borne from y” is such that y and x are respectively the oldest and the second oldest lineages/leaves of the subtree rooted at n. It follows that the given of both the tree and the birth ranking of the lineages resulting from a realization is sufficient to reconstruct its scenario.*

In short, the scenario of a realization fully determines both its tree shape and the birth ranking of its lineages, while the given of both the tree shape and the ranking of a realization determines its scenario. A tree and a birth ranking are *consistent* with one another if there exists a valid scenario corresponding to both of them.

With Remark ??, the number of scenarios leading to a given tree 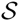 is equal to the number 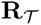 of birth rankings consistent with 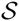. This number depends on the tree considered. It may actually differ between two trees with the same number of leaves.

##### Lemma A1.

*Let* 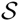 *be a rooted binary labeled tree shape, and* 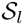 *and* 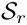 *be the two subtree shapes rooted at the children of its root. A birth ranking* **r** *of the lineages of* 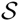 *is consistent with* 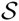 *if and only if:*

1. *the two oldest lineages of* 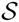 *are the oldest lineage of* 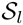; *and of* 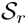,
2. **r**^(^*^l^*^)^, *the restriction of* **r** *to the lineages of* 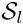, *is consistent with* 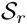,
3. **r**^(^*^r^*^)^, *the restriction of* **r** *to the lineages of* 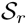, *is consistent with* 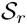.

###### Proof

Let us first assume that r is consistent with 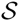. There exists a scenario *E* leading to 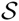 in which the *i*^th^ event is the birth of the lineage of rank (*i* + 1). In particular, its first event is the birth of the second oldest lineage of 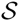 (the oldest one starts the process). The first event corresponds to the root node of 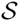, which thus involves the two oldest lineages and splits 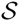 into the subtree containing the second oldest lineage and all its descendants and the subtree containing the oldest lineage and all its descendants except that of the second oldest one and the second oldest one itself. It follows that the two oldest lineages of 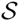 are the oldest lineage of 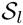 and the oldest one of 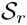. Let *E*^(^*^l^*^)^ be the scenario obtained from *E* by discarding its first event and all the events not involving a lineage of 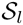. Basically the tree 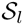 follows the sequence of events of *E*^(^*^l^*^)^ and the corresponding birth ranking is the restriction of **r** to the lineages of 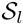. The same holds for 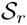.

Reciprocally, let **r**^(^*^l^*^)^ and **r**^(^*^r^*^)^, two birth rankings consistent with 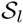 and 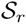 respectively and **r** be obtained by merging **r**^(^*^l^*^)^ and **r**^(^*^r^*^)^ in such a way that the two first lineages of **r** are choosen among 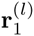 and 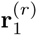. There exist two scenarios *E*^(^*^l^*^)^ and *E*^(^*^r^*^)^ leading to the pair 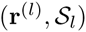 and the pair 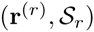 respectively. Let now *E* be the scenario where the first event is “**r**_2_ borne from **r**_1_” and, for all *i* > 1, the event *E_i_* is the birth event of the lineage **r***_i_*_+1_ which belongs either to *E*^(^*^l^*^)^ or to *E*^(^*^r^*^)^. Since the scenarios *E*^(^*^l^*^)^ and *E*^(^*^r^*^)^ are valid, *E* is valid and determines both the tree 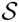 and the birth ranking **r**.

##### Theorem A1.

*The number of birth rankings consistent with a rooted binary labeled tree shape* 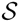 *is*

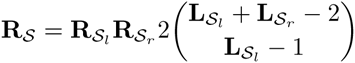

*where* 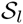, 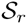, 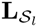 *and* 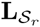 *are the two subtree shapes rooted at the children of the root of* 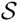 *and their numbers of leaves/lineages respectively.*

*Proof.* From Lemma ??, there are as many rankings consistent with 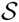 as ways of merging a ranking of 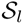 with one of 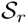, by taking the two first lineages among 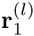 and 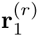. There are:

- 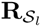 rankings consistent with 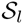,
- 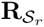 rankings consistent with 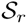,
- 2 ways of setting the two first lineages of a ranking of 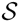 among 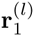 and 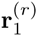,
- 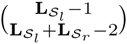 ways of merging the lineages of **r**^(^*^l^*^)^ and **r**^(^*^r^*^)^ except for the two oldest ones (such a merging in fully determined by the ranks occupied by the lineages of **r**^(^*^l^*^)^).

All these possibilities may be combined independently to give a ranking consistent with 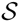.

Since the number of rankings consistent with the tree made of a single lineage is 1, Theorem ?? provides a recursive way to compute 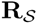 for any tree shape 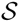.

#### Probability of a tree shape given its number of leaves

Since the labeling of the leaves/lineages is arbitrary (i.e. depends neither on the tree shape nor on their birth ranks), the following remark follows by symmetry.

##### Remark A2.

*In a realization with n lineages arbitrary labeled, all the birth rankings of the (labeled) lineages have equal probability. Since there are n*! *possible rankings, this probability is* 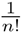.

Until here, we made no assumptions about the realizations or about the processes leading to tree shapes. From now on, we consider only tree shapes arising from pure-birth realizations (i.e. realizations of general birth and death processes in which no death occurs). Moreover, we focus on a large class of processes, which contains the usual diversification models. A process is said *lineage-homogeneous* if, at each event time, all the lineages give birth at a same rate. Such models are called *ERM models* in (Ford *et al*., 2009).

##### Lemma A2.

*Being given the birth ranking of a pure-birth realization of a lineage-homogeneous process, all the tree shapes have probability* 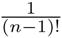.

###### Proof.

Since the realization contains no death and the process is lineage-homogeneous, the *i*^th^ lineage is borne from any of the (*i* – 1) lineages alive at its birth date with equal probability 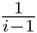, independently of the other events. It follows that, being given the birth ranking, the joint probability of the parenthood of all the lineages is 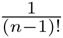.

##### Theorem A.2.

*A tree shape* 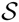 *resulting from a pure-birth realization of a lineage-homogeneous process has probability*

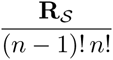

*conditioned on having n leaves.*

###### Proof

From Remark ?? and Lemma ??, the joint probability of a pair tree/ranking is 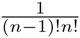. To obtain the probability of a tree 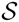 with *n* leaves, we just have to sum these joint probabilities over all the rankings consistent with 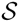, which gives us the result.

### B Complexity index of a tree

The complexity of Algorithm 1 relies on the number of possible before/after assignments of the basic trees encountered during its execution. Let us put 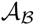 for the number of before/after assignments of a basic tree 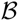 with regard to a time between those of the origin and of the oldest leaf of 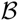. Let 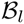 and 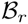 be the subtrees pending to the children of the root of 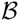. Any before/after assignment of 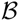 in which the root is set to “before”, is obtained in a unique way by combining an assignment of 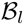 with one of 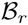. There is only one possible assignment of 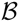 in which the root is set to “after”. It follows that we have

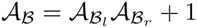

The number of before/after assignments of a basic tree is recursively computed (the tree made of a single leaf has a unique before/after assignment).

The *complexity index* of a tree 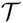 is mainly obtained by summing the number of possible before/after assignments of all the basic trees which have to be considered to compute the likelihood of 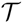. For technical reasons, we actually consider an additional term which is very similar to the number of assignments. Though it certainly be improved, the complexity index predicts quite well the duration of a likelihood computation (see the help of *Diversification,* https://github.com/gilles-didier/Diversification).

### C Sampling extant taxa

Following (Stadler, 2010), we assume here that each extant taxa is independently discovered (or sampled) at contemporary time with a certain probability. Let us define **P***_ρ_*(*n*, *t*) as the likelihood of sampling *n* > 0 lineages at time *t* with the probability *ρ*, by starting from a single lineage at time 0 without fossil meanwhile. The probabilities **P***_ρ_*(0, *t*) and **P***_ρ_*(1, *t*) were already provided in (Stadler, 2010).

For all positive integers n, we have

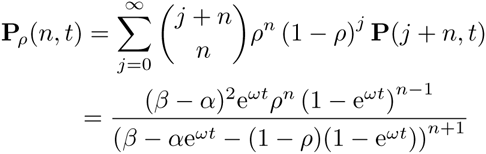

The probability of sampling no lineage at *t*, still by starting from a single lineage at time 0 without fossil meanwhile, is

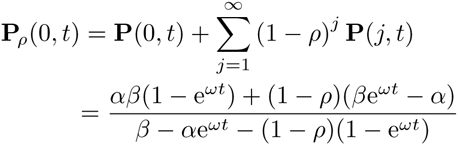

We verified that **P***_ρ_*(0, *t*) and **P***_ρ_*(1, *t*) are well equal to *p*_0_(*t*) and *p*_1_ (*t*) of Theorem 3.1 in (Stadler, 2010) which refer to the same probabilities but which are computed and expressed in a slightly different way.

The likelihood of a reconstructed tree with fossils with extant taxa sampled with the probability *ρ* may be computed in a similar way as under the assumption that all the extant taxa are known. One just needs to replace the probabilities of the form **P**(*n*, *T* – *t*) by **P***_ρ_*(*n*, *T* – *t*) in the calculus. Further tests have to be carried out to check in what extent the sampling probability influences the estimation of the diversification and fossilization rates and in what extent it can be estimated itself.

### D Proportion of lineages unobservable from the fossil record

Let us put **P**_⊘_ for the probability for a lineage to leave no fossil, neither itself, nor its descendant. By assuming that the diversification does not end (i.e. that we are arbitrarily far from the contemporary time), we have that

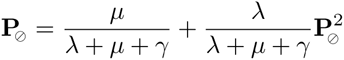

The first term of the right-expression is the probability that the first event occurring on the lineage next its birth is an extinction. The second one is the probability that this event is a speciation giving birth to two lineages leaving no fossil themselves.

The preceding equation can be written as

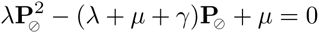
and was already considered in Section 3.2 and (Didier *et al*., 2012).

If *λ* = 0, the unique solution is 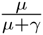. Otherwise, its roots are

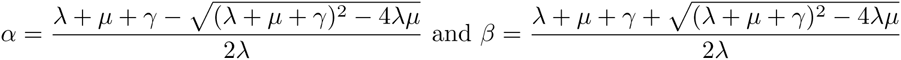

By noting that

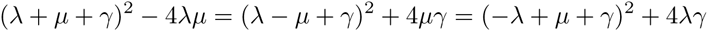
it comes that *α* and *β* are real numbers with *β* ≥ 1 and 0 ≤ *α* ≤ 1.

If *γ* > 0 then *β* > 1 and we have necessarily **P**_⊘_ = *α*, that gives us a natural interpretation of the coefficient *α*. The probability **P**_⊘_ can be itself interpreted as the asymptotical proportion of lineages unobservable from the fossil record. It does not take into account the lineages observable from the contemporary time only.

Remark that **P**_⊘_ is not the complementary probability for species to be fossilized as itself (i.e. before cladogenesis or extinction). This last one, again under the assumption that the diversification does not end, is 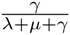.

